## Supplementary information for "Type I interferon shapes brain distribution and tropism of tick-borne flavivirus"

^#^ Contributed equally

**This PDF file includes:**

Figures and legends of figures S1-5

Legends for Video S1-3

Legends for Table S1-2

Table S3

**Other supplementary materials for this manuscript include the following:**

Video S1-3

Table S1-2

Source Data file

**Supplementary figures**
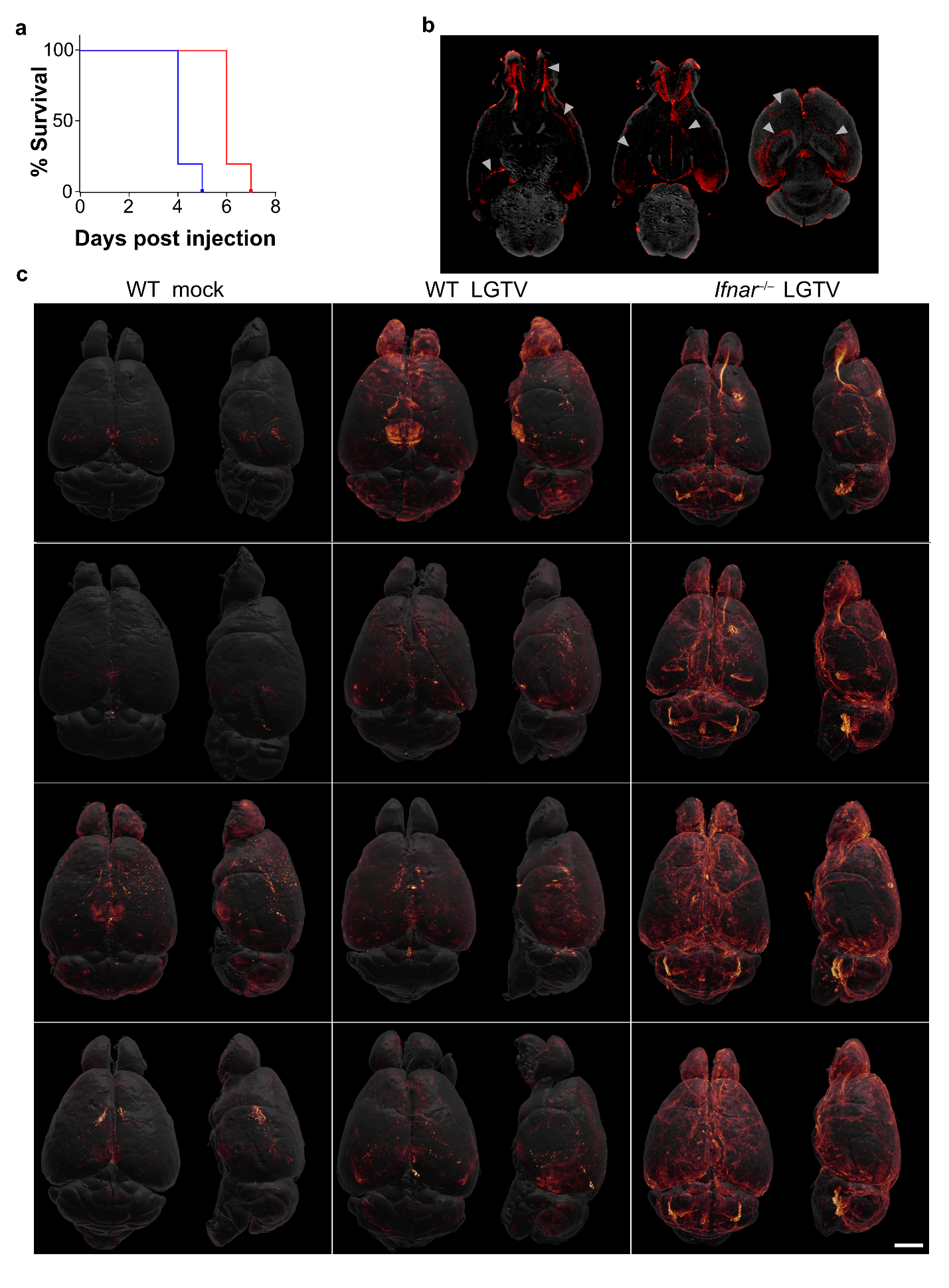

**Supplementary Fig. 1. IFN-I response influences host survival and distribution of LGTV infection. a**  Survival analysis of WT and *Ifnar*^–/–^ mice, intracranially injected with 1000 pfu of LGTV (n = 5). Survival differences between groups were significant (*p* < 0.005, log-rank test). **b** OPT cross sections of a representative *Ifnar*^-/-^ brain showing antibody penetration in the deeper areas of the brain (arrows). **c** OPT-scanned immunolabeled brain reveals the distribution of LGTV infection in the adult mouse brain. Volumetric 3D render of supplementary OPT scans of the brain from mock and LGTV infected mice immunolabeled with antibodies against viral NS5. The signal intensity was normalized within an individual brain and adjusted to identical minimum and maximum. The viral signal was overlaid onto the anatomical outlines created from iso-surface rendering of the autofluorescence signal of each brain. For each image pair, the top and lateral views of the same specimen are shown. Scale bar = 2000 µm.

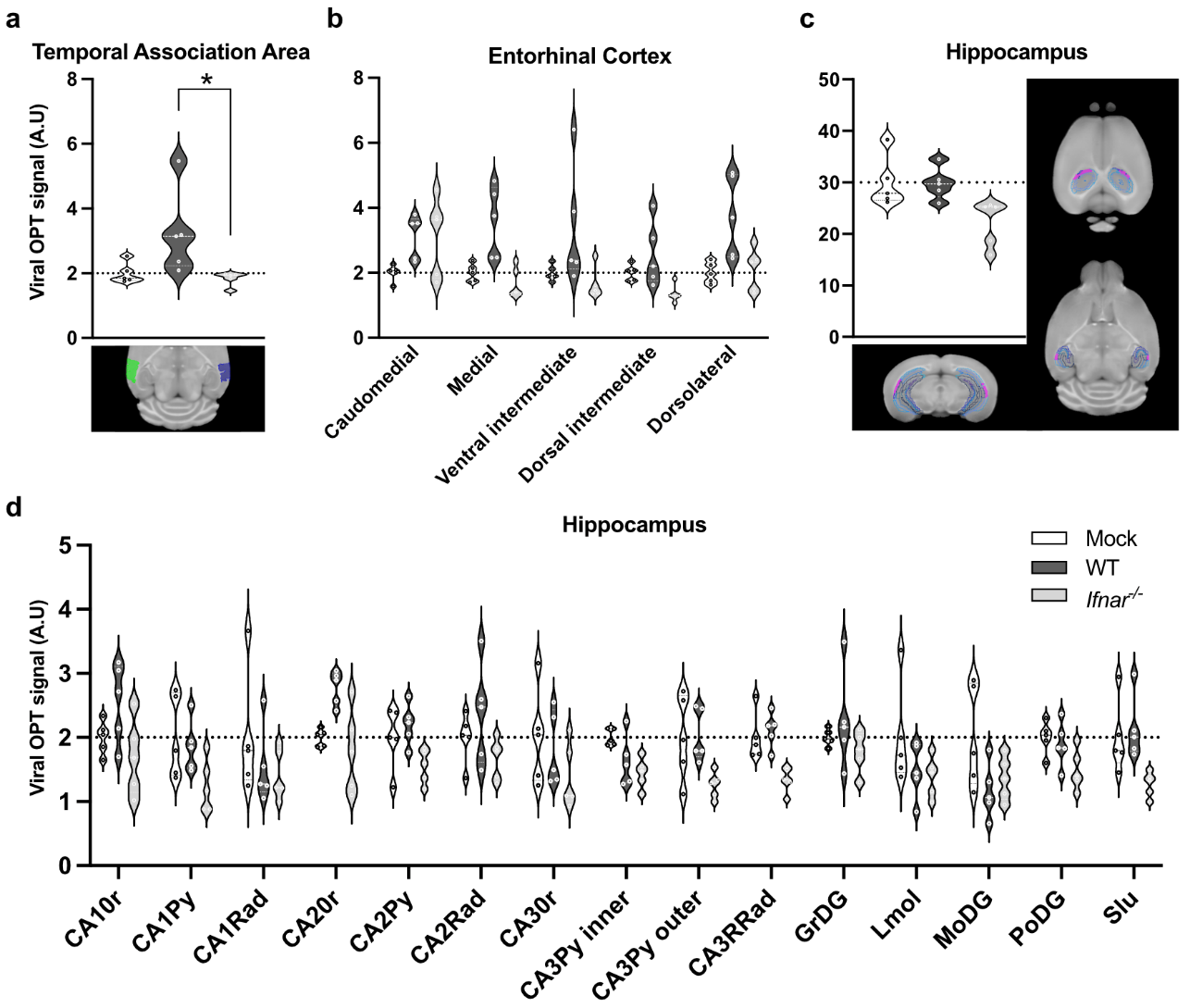

**Supplementary Fig. 2. Image and quantification of viral infection in cerebral cortex regions.** Quantification of viral OPT signal in **a** temporal association area, **b** individual VOIs in the composite entorhinal cortical region, **c** composite hippocampal region, and **d** individual VOIs in the hippocampal region.* p=0.05

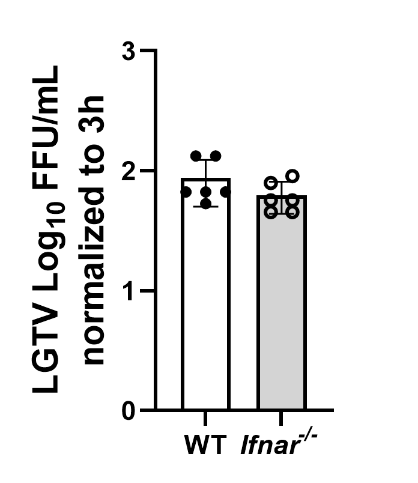

**Supplementary fig. 3 Primary microglia infected with LGTV *in vitro***. Primary microglia were isolated from WT and *Ifnar*^–/–^ brains. The cells were infected with LGTV (MOI 1) for 72 h and viral growth was measured by focus forming assay (n = 5).

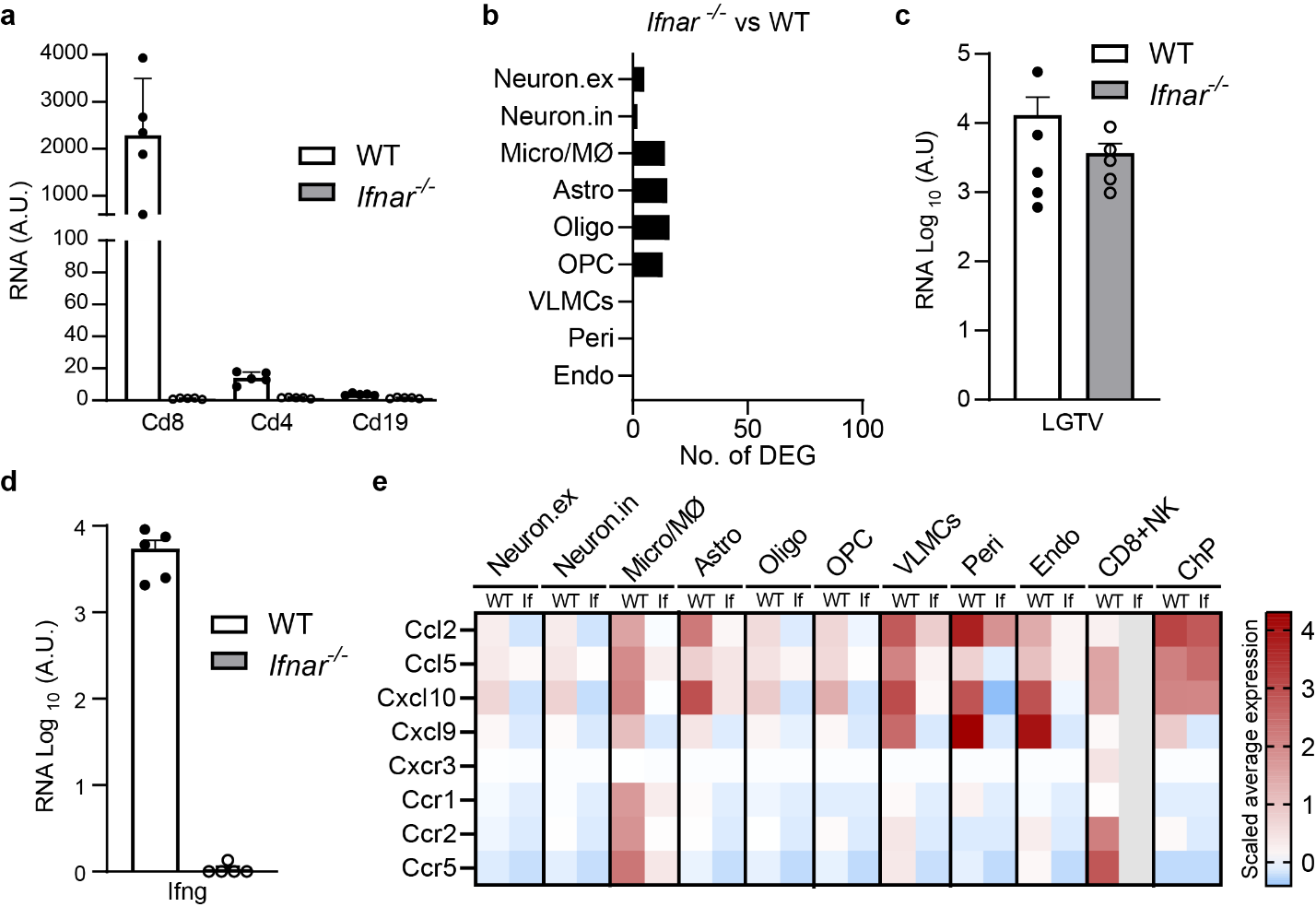

**Supplementary Fig. 4. Transcriptional differences and viral load in WT and *Ifnar^–/–^* mice. a** Gene expression of *Cd8*, *Cd4* and *Cd19* in cortex of infected WT (n=5) and *Ifnar^–/–^* (n=5) mice. Quantified by qPCR and normalized to housekeeping gene *Gapdh*. **b** Number of DEGs (log2FC>1, padj < 0.05) between *Ifnar^–/–^* and WT mice in uninfected samples. **c** Viral load in cerebral cortex at the humane endpoint in WT (n=5) and *Ifnar^–/–^* mice (n=5) infected with 10 000 PFU intracranially. Quantified by qPCR and normalized to housekeeping gene *Gapdh*. **d** Gene expression of *Cd8*, *Cd4* and *Cd19* in cortex of infected WT (n=5) and *Ifnar^–/–^* (n=5) mice. Quantified by qPCR and normalized to housekeeping gene *Gapdh*. **e** Heatmap showing expression of T cell chemoattractants and corresponding receptors upon infection in WT and *Ifnar^–/–^* (If) mice.

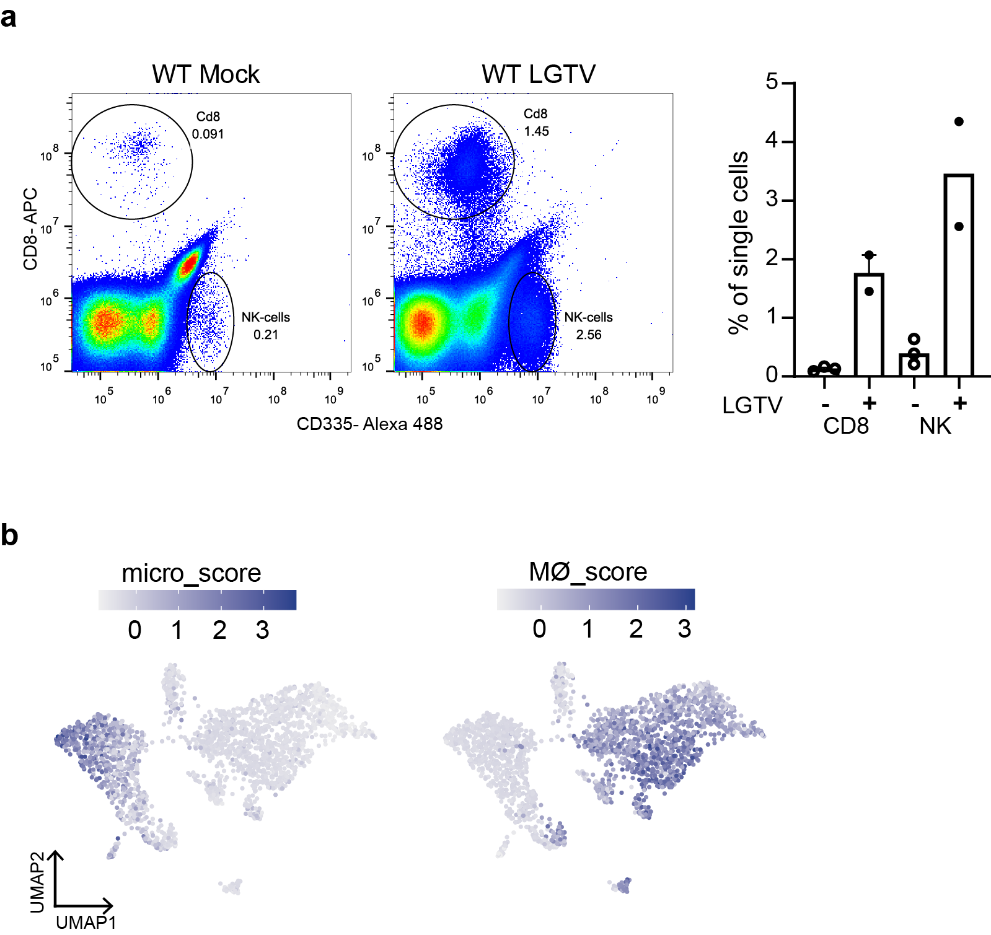

**Supplementary Fig. 5. Infiltration of immune cells in WT and *Ifnar^–/–^* mice. a** Flow cytometry of brain leukocytes from the cerebral cortex of untreated and infected WT mice. Representative scatterplots and percentage CD8+ T cells and NK cells (CD335^+^). **b** Subclustering of 2,363 nuclei of micro/MØ cell subset belonging to all datasets, colored by gene signatures for microglia (*Cx3cr1, P2ry12, Slco2b1, Tmem119*) or MØ (*Slfn4, Ms4a8a, Clec4e, Itga4*) (PMID: 31325960).

**Supplementary Movie 1. 3D renderings of OPT-scanned immunolabeled mouse brain showing the distribution of LGTV infection.** The brains from LGTV infected mice were harvested at endpoint, immunolabeled with anti-NS5 antibody (red glow: normalized to the virus signal intensity within an individual brain) and imaged using OPT.

**Supplementary Movie 2. LSFM showing high-resolution image of the fourth ventricle ChP infected with LGTV.**

(A) Orientation of the brain in the LSFM during image acquisition. The illustration was created with BioRender.com.

(B) Volumetric 3D render of LSFM of LGTV infected fourth ventricle ChP, immunolabeled with anti-NS5 antibodies (red glow). ChP was imaged in 3 tiles at 2.5× magnification and stitched together with 20% overlap. Z = 2000 μm; scale bar = 300 μm. Solid-line square indicates the location imaged in (C). Dashed-line square shows the orientation of (C).

(C) Tomographic section of (B) viewed from the YZ plane at 4× magnification. z = 600 μm.

**Supplementary Movie 3. FIB-SEM volume image of LGTV infected ChP.** Slice-through view of the FIB-SEM volume image with 3D segmented representation of the volume. Segmentation includes replication complexes (yellow), ER (blue), mitochondria (green), and plasma membranes (pale and dark orange).

**Supplementary Table 1. Quantification of OPT signal**

**Supplementary Table. 2. Lists of differentially expressed genes, Reactome pathways and cell-cell communication.**

**Supplementary Table 3. Key Antibody Table**

| Antibodies | Origin | Clonality | Working dilution | Use in the current study | Company (Cat#) |
| --- | --- | --- | --- | --- | --- |
| Anti-chicken Alexa Fluor 488 | Goat | Polyclonal | 1:400 | IHC | Thermo Fisher Scientific (A11039) |
| Anti-chicken Alexa Fluor 555 | Goat | Polyclonal | 1:500 | IHC | Thermo Fisher Scientific (A21437) |
| Anti-chicken Alexa Fluor 680 | Goat | Polyclonal | 1:500 | OPT, IHC | Abcam (ab175779) |
| Anti-mouse Alexa Fluor 647 | Goat | Polyclonal | 1:500 | IHC | Abcam (ab150115) |
| Anti-rabbit Alexa Fluor 488 | Donkey | Polyclonal | 1:500 | IHC | Thermo Fisher Scientific (A21206) |
| Anti-rabbit Alexa Fluor 594 (pre-absorbed) | Donkey | Polyclonal | 1:500 | OPT | Abcam (ab150064) |
| Anti-mouse CD45 Alexa 488 | Rat | Monoclonal | 1:100 | IHC | Invitrogen (53-0451-82) |
| AQPI | Mouse | Monoclonal | 1:50 | IHC | Santa Cruz Biotechnology (sc-32737) |
| Calb | Rabbit | Monoclonal | 1:500 | IHC | Abcam (ab108404) |
| CD8 | Rat | Monoclonal | 1:200 | IHC | BD Pharmigen (561093) |
| CD335 | Rat | Monoclonal | 1:500 | IHC | Thermo Fisher Scientific (AB_1724164 |
| DCX | Rabbit | Polyclonal | 1:800 | IHC | Cell Signaling (4604) |
| Iba1 | Rabbit | Polyclonal | 1:400 | IHC | Histolab (CP290) |
| NS5* | Chicken | Polyclonal | 1:1000 | OPT, IHC | Agrisera AB |
| TMEM119 | Rabbit |  | 0.5µg/ml | IHC | Abcam (ab209064) |
| TUBB3 | Rabbit | Polyclonal | 1:3000 | IHC | BioLegend (PRB-435P) |

*Affinity-purified NS5 antibody was produced in chicken, according to the manufacturer’s protocol, using the following peptide sequence of NS5 from tick-borne encephalitis virus strain Torö (GenBank: DQ401140): (carboxylated)-CMDRHDLHWELRLESS-(amidated).
